# Hippocampal spatio-temporal cognitive maps adaptively guide reward generalization

**DOI:** 10.1101/2021.10.22.465012

**Authors:** Mona M. Garvert, Tankred Saanum, Eric Schulz, Nicolas W. Schuck, Christian F. Doeller

## Abstract

The brain forms cognitive maps of relational knowledge, an organizing principle thought to underlie our ability to generalize and make inferences. However, how can a relevant map be selected in situations where a stimulus is embedded in multiple relational structures? Here, we find that both spatial and temporal cognitive maps influence generalization in a choice task, where spatial location determines reward magnitude. Mirroring behavior, the hippocampus not only builds a map of spatial relationships but also encodes temporal distances. As the task progresses, participants’ choices become more influenced by spatial relationships, reflected in a strengthening of the spatial and a weakening of the temporal map. This change is driven by orbitofrontal cortex, which represents the evidence that an observed outcome is generated from the spatial rather than the temporal map and updates hippocampal representations accordingly. Taken together, this demonstrates how hippocampal cognitive maps are used and updated flexibly for inference.

## Introduction

As humans we live in complex, ever-changing environments that often require us to select appropriate behaviors in situations never faced before. Luckily, our environment is replete with statistical structure and our experiences are rarely isolated events^1^. This allows us to predict outcomes that were never experienced directly by generalizing information acquired about one state of the environment to related ones^2^. Indeed, humans and other animals generalize across spatially or perceptually similar stimuli^3–6^ as well as across stimuli forming associative structures such as those acquired in a sensory preconditioning task^7, 8^. Generalization also occurs in reinforcement learning tasks where the same latent state determines the outcome associated with choosing different stimuli^9, 10^.

For generalization to be possible, an appropriate neural representation of stimulus relationships is required. Many studies have shown that spatial relationships, such as distances between landmarks, are represented in a hippocampal cognitive map^11, 12^, which enables flexible goal-directed behavior beyond simple stimulus-response learning^13^. More recently, it has been suggested that the same organizing principle might also underlie the representation of relationships between non-spatial states such as perceptual^14–19^ or temporal relationships between stimuli^20–22^, or associative links between objects^23–26^. Interestingly, cognitive maps even form incidentally and in the absence of conscious awareness^23^. This suggests that the hippocampus automatically extracts the embedding of a stimulus in multiple relational structures^27^, even for stimulus features that are not directly task-relevant^28^.

If stimuli are part of multiple relational structures such as space and time, this raises the question how the representation that is most beneficial for reward maximisation and generalization can be selected^29^. One region implicated in this process is the orbitofrontal cortex (OFC), known to represent task states in situations where these are not directly observable^24, 30^. Little is known, however, about how information in the OFC about the task-relevance of different maps relates to corresponding changes in the representation of cognitive maps in the hippocampus^31, 32^.

Here, we combined virtual reality with computational modeling and functional magnetic resonance imaging (fMRI) to show that participants represent spatial as well as temporal stimulus relationships in hippocampal maps. The degree to which each map was represented neurally determined the degree to which it was used for generalization in a subsequent choice task, even though only the spatial location determined the magnitude of rewards. Notably, the neural representation of each map and its influence on choice changed over the course of the choice task through an OFC signal reflecting the evidence that the spatial rather than the temporal map caused the observed outcome. Together, our results provide a computational and neural mechanism for the representation and adaptive selection of hippocampal cognitive maps during choice.

## Results

### Participants used knowledge about stimulus relationships to generalize value

To examine how humans use information about stimulus relationships for generalization and inference, forty-eight healthy human participants (mean age 26.8 ± 3.8 years, 20 − 34 years old, 27 male) took part in a 3-day experiment that involved first learning to locate 12 monster stimuli in a virtual arena, followed by a choice task in which spatial knowledge could be used for predicting rewards (Figure 1a).

**Figure 1.**
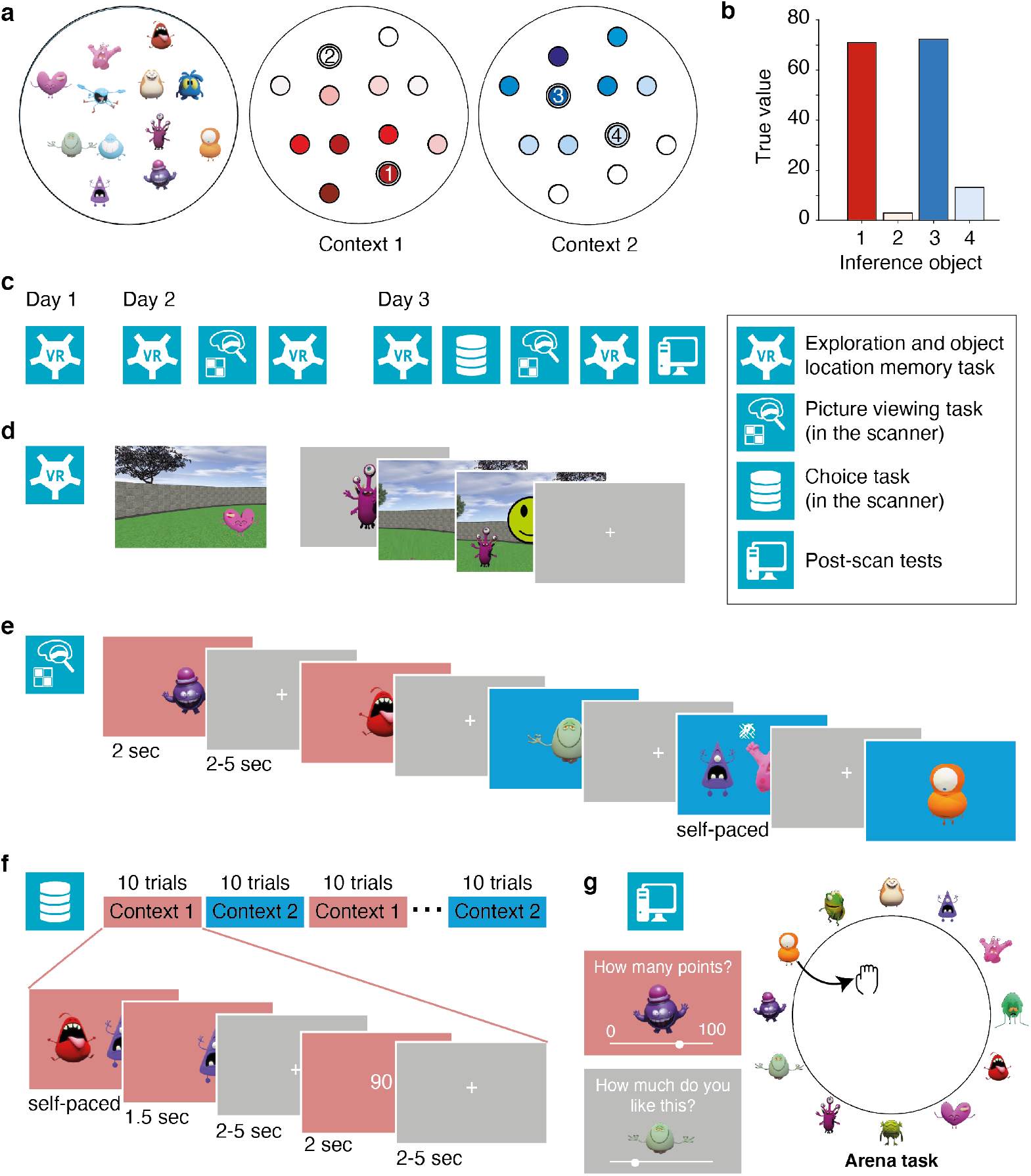
Experimental design. **a** Spatial position of monsters during the navigation tasks and value distribution associated with the same monsters in context 1 and 2 in the choice task. Darker colors indicate higher values. Numbered circles indicate the location of inference objects that were never presented during the choice task. **b** True values of the four inference objects. **c** Tasks performed on the three subsequent days, see text. **d** Exploration and object location memory tasks. In the exploration task, participants navigated around a virtual arena with button presses corresponding to forward, backward, right and left movements. Monsters appeared when they were approached, but were never all visible at the same time. In the object location memory task, participants were instructed to navigate to the position of a cued monster (each monster cued once in each block). Feedback indicated how far away the positioned object was from the correct object location. On day 1, participants performed between five and ten blocks (depending on performance) of the exploration and the object location memory task in alternation. On subsequent days, only one block of the object location memory task was performed before and after scanning without feedback. **e** Picture viewing task performed in the scanner. Participants were presented with monsters one after another. When two monsters appeared, participants were instructed to choose the monster that was closer in space to the preceding monster (map symbol) or the monster that was more similar in value to the preceding monster (coins symbol, day 3 only). On day 2, the background color was irrelevant for the task, on day 3 it indicated the context determining the stimulus values. **f** Choice task performed in the scanner. Participants were instructed to maximize accumulated points by choosing the monster associated with a higher reward. Participants were told that the monsters had different values in two different contexts, and that the relevant context was signalled by the background color. The values associated with each monster in the two contexts were learned in alternation, with ten blocks of context 1 followed by ten blocks of context 2, and so forth. **g** At the end of day 3, four post-tests were performed: Participants indicated for each monster how many points they would receive in each of the two contexts and how much they liked each monster. They were then asked to arrange the monsters in terms of their similarity in a circle in such a way that monsters that were considered similar were positioned near each other (Arena task 1). Lastly, participants were instructed to imagine a top-down view of the arena they had navigated around and to place the monsters in the corresponding location (Arena task 2).

On day 1, participants performed multiple exploration blocks in which they were instructed to remember the location of the stimuli while freely navigating in the arena (Figure 1c, d). Stimuli became visible when they were approached, but were otherwise invisible. Exploration policies differed substantially between individuals (Figure 2a, Supplementary Figure S1). As a result, participants experienced different temporal relations between the monsters, which could also deviate from the spatial distances between objects. For example, some participants visited objects in a stereotyped order, whereas others navigated mostly around the border of the arena or systematically scanned the environment from top to bottom (Figure 2a).

**Figure 2.**
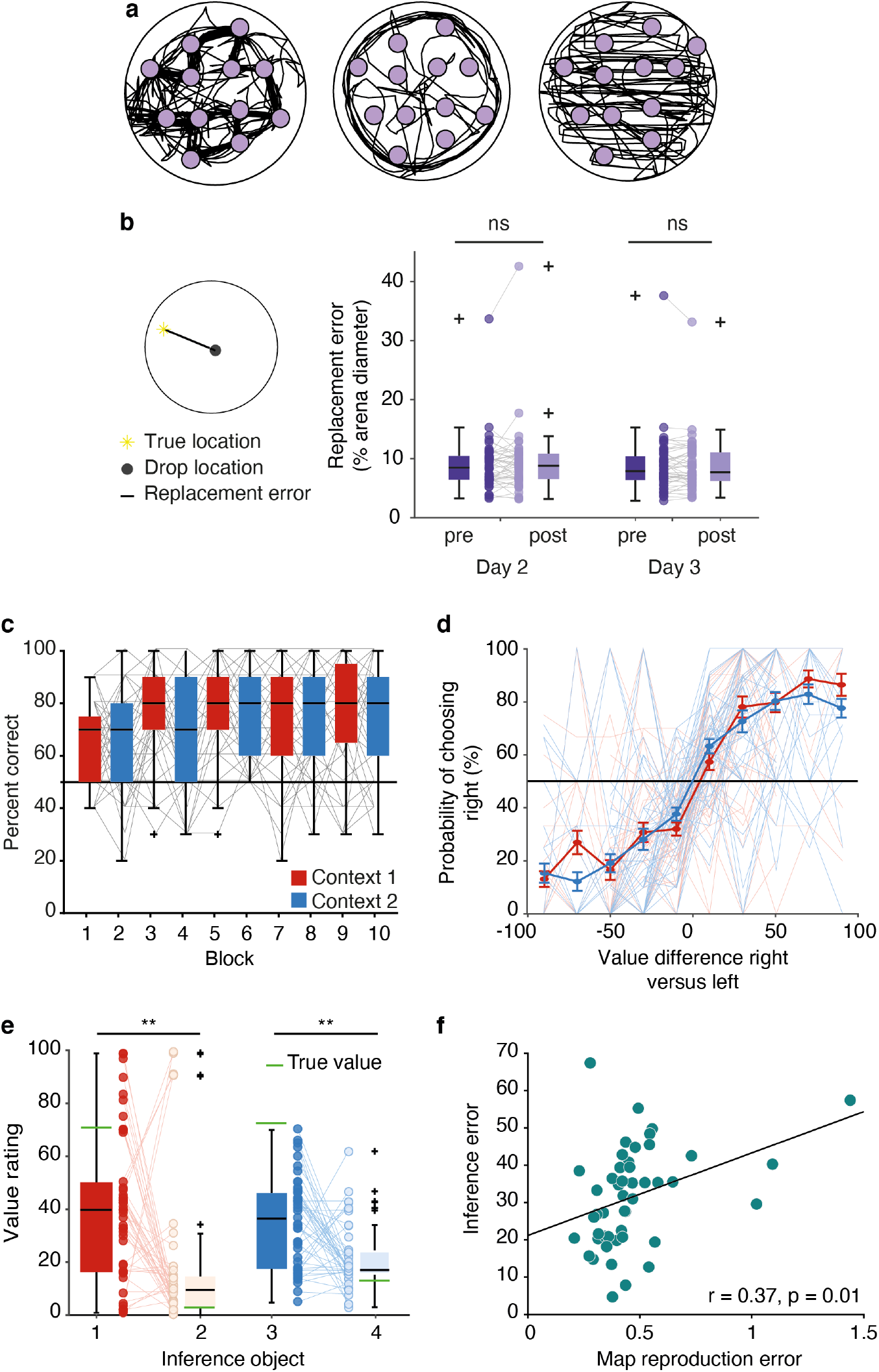
Behavioral results. **a** Trajectories of three example participants during the exploration phase on day 1. Purple dots indicate the stimulus locations and black lines the participant trajectories. See all participants’ trajectories in Supplementary Figure S1. **b** replacement error for days 2 and 3, before (pre) and after (post) the scanning session. The replacement error was defined as the Euclidean distance between the true location and the drop location. The replacement error did not differ significantly between those four sessions (all *p* > 0.05), see object positioning at the end of the learning phase on day 1 in Supplementary Figure S2. **c** Percent correct of choices over the course of the choice task. Trials are divided into ten sub-blocks of ten trials each with a constant context. **d** Probability of choosing the right option as a function of the difference in value between the right and the left option, separately for each context. **e** Value rating for the inference stimuli at the end of the study. **f** Correlation between the the map reproduction error (root-mean-square error between the true *z*-scored spatial distances and the *z*-scored distances in the arena task) and the root-mean-square error for the inference ratings. Data in **b, c** and **e** are plotted as group-level whisker-boxplots (center line, median; box, 25th to 75th percentiles; whiskers, 1.5 × interquartile range; crosses, outliers). Error bars in **d** denote standard error of the mean. Circles and transparent lines in **b**-**f** represent individual participant data. ** *p* < 0.01, ns = not significant

After each exploration block, participants performed an object location memory task. Participants were first teleported to a random location in the arena and instructed to then navigate to the hidden location of a presented object. Feedback indicated how far away the current position was from the correct stimulus location. The session terminated when the replacement error averaged across all monsters in a block was below 3*vm* (vm = virtual meter; 3vm correspond to < 10% of the arena’s diameter) and at least five and at most ten blocks had been completed. At the end of the learning phase, participants could position the stimuli in the correct location (Supplementary Figure S2a). Before and after each imaging session on days 2 and 3, participants also performed one block of the object location memory task without feedback. The replacement error did not differ between sessions (Figure 2b), confirming that no new learning took place. In a spatial arena task at the end of the 3-day study, participants also accurately reproduced the stimulus arrangement when instructed to drag-and-drop stimuli imagining a top-down view on the spatial arena (Figure 1g). Participants thus learned the spatial arrangement of the stimuli well.

In a choice task performed in the MRI scanner on day 3, participants were presented with two stimuli simultaneously and instructed to select the one that was associated with a higher reward (Figure 1f). Participants were told that the reward magnitude was determined by the stimulus location in space (Figure 1a). Participants did initially not know which locations were rewarding, but they could combine their knowledge about the stimulus relationships with previously experienced reward contingencies to infer the rewards of stimuli they had not yet experienced. In order to decorrelate spatial distance and reward relationships, we introduced two contexts with different reward distributions (Figure 1a). Participants performed alternating choice blocks for each context, with the context signaled by the background color. Participants learned to perform the task rapidly (Figure 2c) and their choices were a function of the difference in value between the stimuli presented on the left and the right on the screen in both contexts (Figure 2d, context 1: *t*(47) = 10.0, *p* < 0.001, context 2: *t*(47) = 12.1, *p* < 0.001).

To test whether participants could use their knowledge about the stimulus relationships to generalize, two stimuli per context were never presented during the choice task (“inference stimuli”, Figure 1a, b). A value rating at the end of the study (Figure 1g) showed that participants were able infer which of the two inference stimuli had a higher value in each context (Figure 2e; repeated measures ANOVA, *F*(1, 46) = 21.4, *p* < 0.001), reflecting that they combined their knowledge about the stimulus location with knowledge about associated rewards of nearby stimuli. The error between the true inference values and the value ratings was larger in participants who struggled to reproduce the spatial map as indicated by a larger error between the true z-scored spatial distances and the z-scored distances in the arena task (“Map reproduction error”, *r* = 0.37, *p* = 0.01, Figure 2f). This demonstrates that participants exploited knowledge about stimulus relationships to infer unseen values.

### Cognitive maps of spatial and temporal stimulus relationships explain generalization

The fact that participants could successfully infer the values of the inference stimuli suggests that they formed a representation of the stimulus relationships. But stimulus relationships were learned during free exploration, which was typically non-random and differed substantially between participants (Figure 2a, Supplementary Figure S1). This means that the experienced temporal distances between the objects differed meaningfully from their spatial distances in most participants (Supplementary Figures S6 and S4b). Intelligent agents should keep track of both the spatial distance as well as the temporal relationships between objects, since either feature may become relevant for generalization. We therefore reasoned that the brain may extract two relational maps: one reflecting spatial distances between stimuli and the other one reflecting temporal relationships.

To test explicitly to what extent generalization was guided by the spatial or temporal maps – or a combination of both – we fitted Gaussian process (GP) models to participants’ choices (see Online Methods). The GP predicts rewards for a novel stimulus based on the rewards associated with all other stimuli, weighted by their similarity to the novel stimulus. Since the similarity function determines how the GP generalizes, we can express hypotheses about what cognitive map participants use by pairing GPs with similarities implied by spatial or temporal maps.

Specifically, generalizing using a spatial cognitive map corresponds to pairing the GP with a similarity function that decays with Euclidean distance. Generalizing using a temporal cognitive map corresponds to pairing the GP with a similarity function that decays with temporal distance. We constructed these temporal similarities based on individual participants’ navigation runs from day 1: Using their stimulus visitation history from the exploration phase, we computed each participants’ successor representation^33^, reflecting the expected number of visits of any stimulus *s*′ given a starting stimulus *s*. This can be transformed into a probability that two stimuli are visited in direct succession (see Online Methods). We then computed temporal similarities based on the diffusion distance^6, 34^ implied by these transition probabilities (Figure 3a).

**Figure 3.**
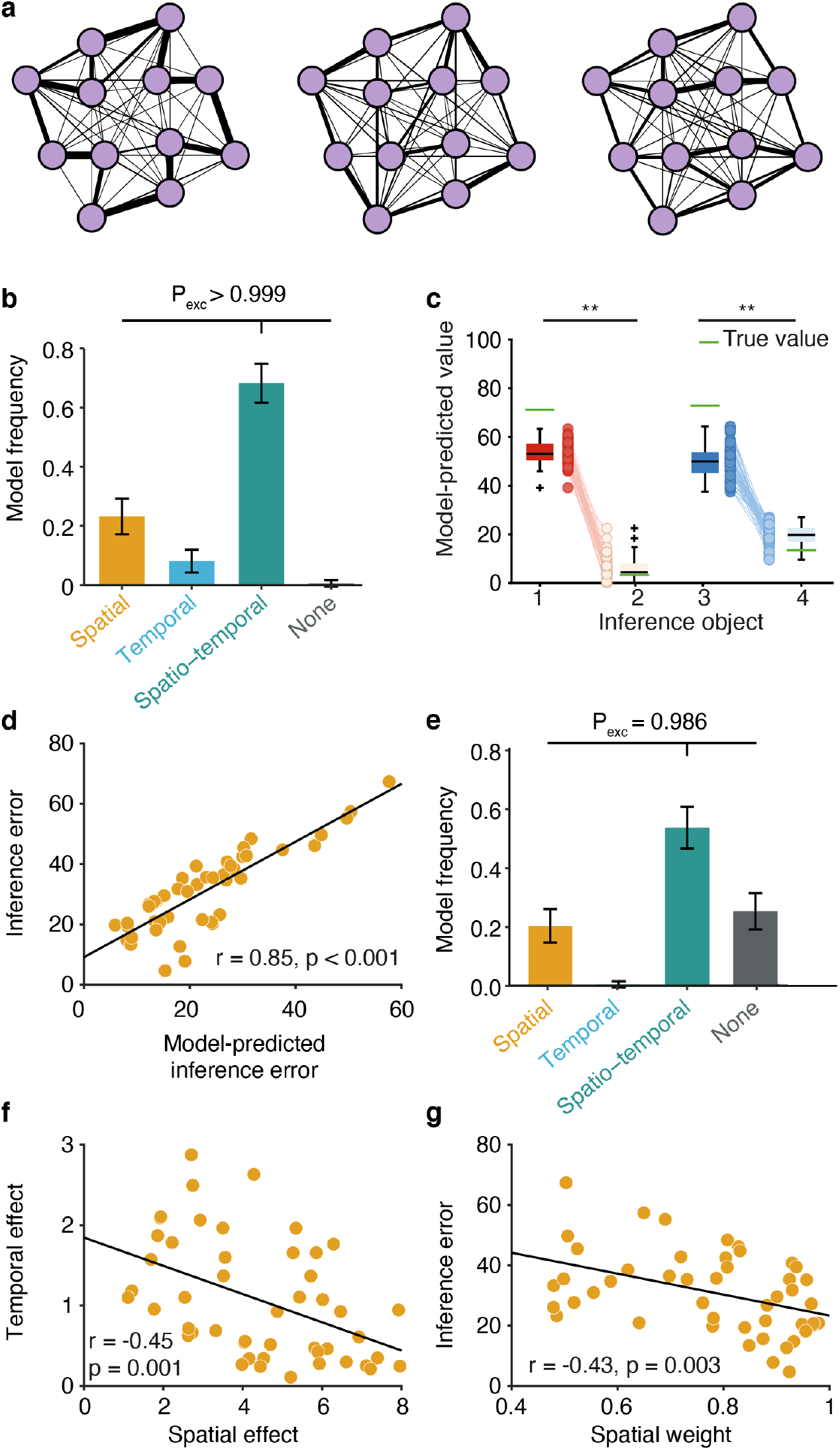
Modeling results suggest that participants generalized over spatial and temporal stimulus relationships. **a** Graph representation corresponding to the three example exploration paths in Figure 2a. **b** Model comparison: Model frequency represents how often a model prevailed in the population. The error bars represent the standard deviation of the estimated Dirichlet distribution. The winning model generalizes values according to a combination of spatial and temporal relationships between stimuli. **c** Inference performance as predicted by the model. Depicted are the inferred values for the inference objects in analogy to the participant ratings in 2e. **d** Relationship between inference error predicted by the model and actual inference error in participants’, value ratings. **e** Model comparison for the value ratings for the inference objects at the end of the study. The winning model generalizes values according to a combination of spatial and temporal relationships between stimuli. **f** Correlation between the spatial and temporal effects on choice behavior. **g** Correlation between the relative spatial weight as estimated by the model and inference error. Data in **c** are plotted as group-level whisker-boxplots (center line, median; box, 25th to 75th percentiles; whiskers, 1.5 × interquartile range; crosses, outliers). Circles and transparent lines represent individual participant data. ** *p* < 0.01, P_exc_ = exceedance probability.

Finally, kernel functions can be added or multiplied together^35^ to model function learning where generalization may be guided by a combination of multiple similarity functions^36, 37^. As such, the hypothesis that both the spatial *and* temporal maps guide generalization together is captured in the spatio-temporal GP, which uses the additive composition of the *spatial* and the *temporal* similarities to generalize.

To test which map best explained how participants generalized rewards, we created three GP models that generalized based on either spatial, temporal or spatio-temporal relationships between monsters. Then, for each trial, we made each GP model predict the reward of both monsters, conditioning the GPs on all monster-reward pairs observed in the relevant context up to that point. We also compared these models to a “mean tracker” model that assumes participants only learn about directly experienced stimulus-reward associations, without generalization (see Online Methods).

To fit our models to participants’ choices, we entered the predicted difference in reward between the two presented monsters in a mixed-effect logistic regression model with random slopes per participant^38^, and determined the maximum likelihood hyper-parameters using grid search. We then computed model frequency based on the leave-one-out cross-validated log-likelihood (leaving one trial out) for each model^39^.

The model generalizing based on the compositional, spatio-temporal similarities explained participants’ choices best (Figure 3b; model frequency = 0.681, XP > 0.999, see Supplementary Figure S3 for full modeling results). This model performed substantially better than the temporal model (model frequency = 0.08), the spatial model (model frequency = 0.23) and the mean tracker (model frequency = 0.005). The model also reproduced the difference in value rating for the high- and the low-inference stimuli (Figure 3c; repeated measures ANOVA, *F*(1, 47) = 2602.3, *p* < 0.001). Across participants, the root-mean-square error between true values and values predicted by the winning model was highly correlated with the root-mean-square error between the true values and the value ratings provided by participants (Figure 3d, *r* = 0.85, *p* < 0.001).

Furthermore, participants’ value ratings for the inference objects at the end of the study were also predicted best by a spatio-temporal model (Figure 2e). This demonstrates that behavior in two independent parts of the study, the choice task and the inference test, was influenced by both spatial and temporal knowledge about stimulus relationships. Notably, the value ratings for the stimuli whose values could be directly sampled were best predicted by the mean tracker model, rather than the spatio-temporal GP (Supplementary Figure S3a). This suggests that participants evoked specific memories of stimulus-reward associations where possible, but relied on the spatio-temporal map when they needed to construct values of stimuli which were not directly experienced (Supplementary Figure S3c).

We estimated effect sizes for the spatial and the temporal component as the participant-specific random effects in a model where the spatial and temporal predictors competed to explain variance in participants’ choices. Spatial weights were defined as the relative contribution of the spatial compared to the temporal predictor. Both the spatial and the temporal relationships had non-zero influence on choice behavior and the effect sizes were negatively correlated (Figure 3f, *r* = − 0.45, *p* = 0.001), suggesting that participants tended to rely predominantly on one of the two maps for guiding choice. Consistent with the fact that the spatial, but not the temporal relationships, were relevant for generalization, participants whose choices were driven more by the spatial relationships compared to the temporal ones performed better in the inference test (Figure 3g, *r* = −0.43, *p* = 0.003).

### Spatial and temporal stimulus relationships represented in the hippocampal system influence choice

Our modeling results suggest that participants generalized values based on both the spatial and temporal relationships experienced between stimuli during the exploration phase. To investigate the neural representation of these relationships, we scanned participants before the choice task on day 2 and after the choice task on day 3 using fMRI. During these imaging sessions, stimuli on the two background colors were presented in random order (Figure 2e). Once after each stimulus on each background color (i.e. in 24 of 144 trials), participants were presented with two stimuli and instructed to either report which one was closer in space or more similar in value in the given context (on day 3 only) to the preceding stimulus. Participants performed this task well above chance (correct performance on day 2: 81 ± 10% (distance judgement); day 3: 78 ± 12% (distance judgement) and 68 ± 14% (value judgement), mean ± standard deviation, all *p* < 0.001).

We used fMRI adaptation^40, 41^ to investigate the representational similarity of the 12 stimuli. This technique uses the amount of suppression or enhancement observed when two stimuli are presented in direct succession as a proxy for the similarity of the underlying neural representations. In line with previous work demonstrating similar effects for graph-like structures^23^, we hypothesized that in regions encoding a cognitive map of the stimulus relationships, the size of the cross-stimulus adaptation effect should scale with spatial or temporal distance between stimuli. Based on previous work, we expected the hippocampal formation to be a candidate region for representing such cognitive maps^23^ and therefore focus on a bilateral region comprising the hippocampus, the entorhinal cortex and the subiculum (see mask used for small-volume correction in Supplementary Figure S5). We tested for adaptation effects by including spatial and temporal distances as parametric modulators in the same general linear model (GLM).

We found a significant cross-stimulus enhancement effect that scaled with spatial distance in session 3 (after the choice task) in the right hippocampal formation (Figure 4a, *t*(47) = 3.86, *p* = 0.045, [24, − 28, − 16]). A cluster in the left hippocampal formation trended in the same direction (*t*(47) = 3.63, *p* = 0.08, [− 12, − 36− 6]). No voxels survived the conservative correction procedure for the temporal relations. One reason for this could be that different participants represented the spatial and temporal aspects to different degrees, with a stronger representation of the spatial map across the group as a whole. Indeed, in most participants (44 out of 48), the spatial component contributed more to generalization during choice than the temporal component (*t*(47) = 9.9, *p* < 0.001). We therefore investigated whether the strength of the neural representation predicted the degree to which an individual was influenced by either spatial or temporal distances in the choice task.

**Figure 4.**
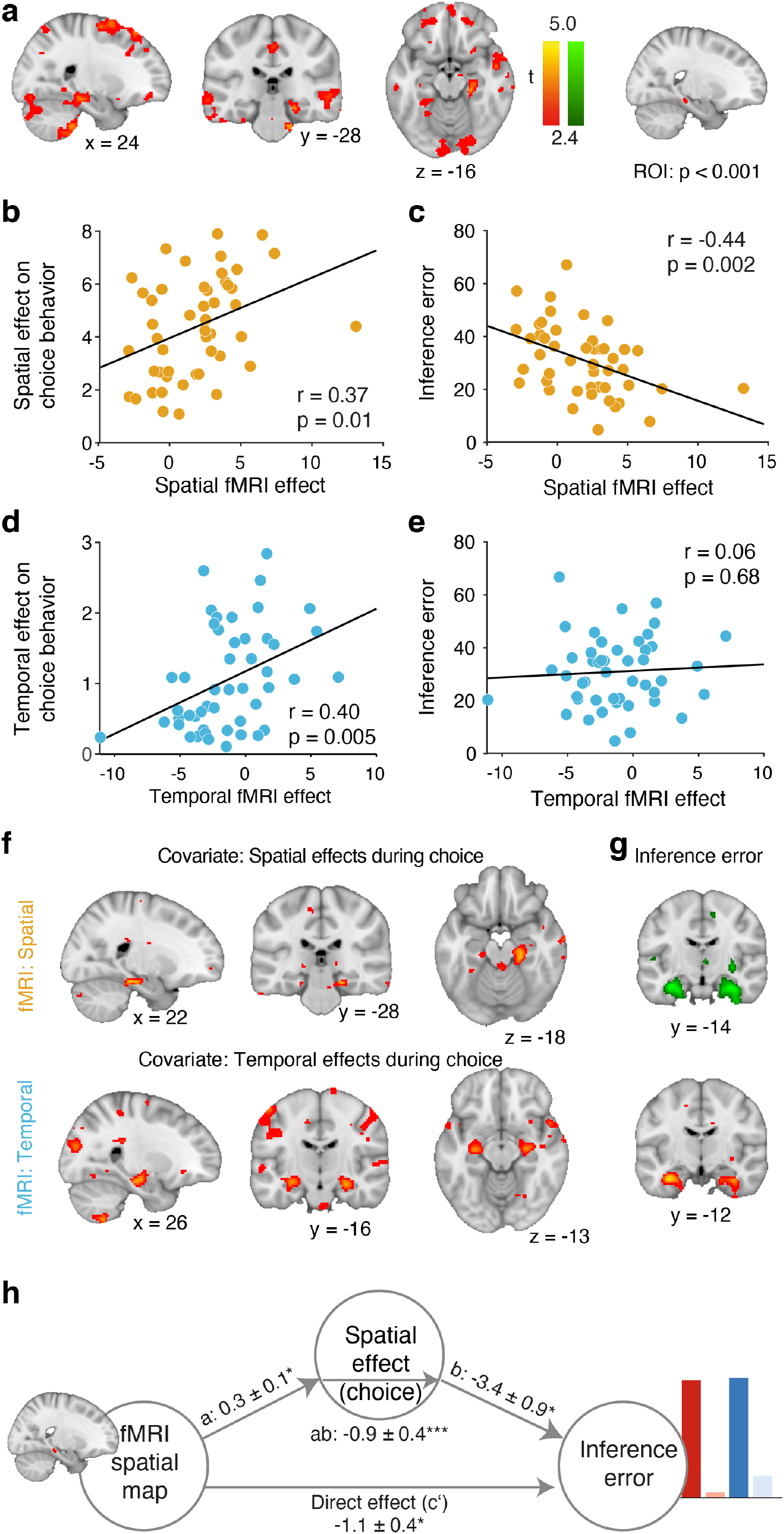
Spatial and temporal cognitive maps in the hippocampal formation are related to generalization and inference. **a** Whole-brain analysis showing a cross-stimulus enhancement effect in the scanning session after the choice task (session 3) that scales with spatial distance. For illustration purposes, voxels thresholded at *p* > .01 (uncorrected) are shown; only the right hippocampal cluster survives correction for multiple comparisons. **b** Correlation between the spatial cross-stimulus enhancement effect extracted from the right hippocampal ROI depicted in **a** (thresholded at *p* < 0.001) and the spatial effects governing decisions in the choice task. **c** Correlation between the spatial cross-stimulus enhancement effect extracted from the right hippocampal ROI depicted in **a** and the root-mean-square error between ratings for the inference stimuli and their true value. **d** Correlation between temporal cross-stimulus enhancement effect extracted from the right hippocampal ROI depicted in **a** and the temporal effects governing decisions in the choice task. **e** Correlation between the temporal cross-stimulus enhancement effect extracted from the right hippocampal ROI depicted in **a** and the root-mean-square error between ratings for the inference stimuli and their true value. **f** Whole-brain analysis where spatial effects (top) and temporal effects (bottom) describing generalization during choice are entered as second-level covariates for the spatial and temporal cross-stimulus enhancement effects. Both analyses reveal significant clusters in the hippocampal formation. **g** Whole-brain analysis where the inference error is entered as second-level covariate for the spatial and temporal cross-stimulus enhancement effects. This analysis reveals a negative effect for the spatial map and a positive effect for the temporal map in the hippocampal formation. **h** Mediation path diagram for inference error as predicted by the hippocampal map and spatial effects. **a, f and g** are thresholded at *p* < 0.01, uncorrected for visualization. ** *p* < 0.01; *** *p* < 0.001

To test this, we first extracted parameter estimates for the spatial and temporal maps from the above-identified region of interest (ROI) in the right hippocampal formation showing a cross-stimulus enhancement effect that scaled with spatial distance (Figure 4a, masking threshold *p* < 0.001). A significant correlation with the spatial and temporal effects on choice behavior confirmed a relationship the neural representation of the respective maps in this region and generalization behavior (Figure 4b, d), spatial: *r* = 0.37, *p* = 0.01, temporal: *r* = 0.40, *p* = 0.005). We also found that the representation of the spatial, but not the temporal map in this ROI can be linked to performance in the later, independent inference test that depended on spatial knowledge (spatial: *r* = − 0.44, *p* = 0.002, temporal: *r* = 0.06, *p* = 0.7, Figure 4c, e).

To investigate whether the relationship between spatial and temporal influences on behavior and neural map representation is specific to the hippocampus, we included spatial and temporal effects on choice behavior as covariates on the second level in the GLM that was used to identify spatial and temporal cross-stimulus enhancement effects above. For both spatial and temporal maps we found precisely localized clusters in the hippocampal formation, where the spatial and temporal fMRI effects were larger the stronger the respective map’s influence on behavior (Figure 4f, spatial: *t*(47) = 4.45, *p* = 0.009, [22, − 28, − 18], temporal: *t*(47) = 4.19, *p* = 0.02, [26, − − 20, − − 28], *t*(47) = 4.14, *p* = 0.02, [28, − 14, − 16] and *t*(47) = 3.91, *p* = 0.04, [− 28, −16, −13]). Furthermore, the representation of the spatial map in the hippocampus was stronger and the representation of the temporal map was weaker in individuals who made smaller inference errors (Figure 4g, spatial: *t*(47) = 5.08, *p* = 0.002, [32, − 14, − 25] and *t*(47) = 4.95, *p* = 0.002, [− 32, − 14, − 22], temporal: *t*(47) = 4.53, *p* = 0.007, [− 32, − 12, − 25]). This suggests that participants who represented the spatial map more strongly in the hippocampal formation also generalized more according to spatial distances in the choice task and performed better in the inference task, with the reverse pattern for the temporal relationships.

To test whether the hippocampal spatial map formally mediated the impact of the neural representation on inference performance, we related the parameter estimates for the spatial map extracted from the right hippocampal ROI to both the spatial effects as estimated from behavior in the choice task as well as the inference performance using single-level mediation^42, 43^. The path model jointly tests the relationship between the neural representation of the spatial map and the degree to which spatial relationships influenced generalization in the choice task (path a), the relationship between spatial weights in the choice task and inference performance (path b), and a formal mediation effect (path ab) that indicates that each explains a part of the inference performance effect while controlling for effects attributable to the other mediator. All three effects were significant (path *a* = 0.26, *SE* = 0.10, *p* = 0.01; path *b* = − 3.40, *SE* = 0.92, *p* = 0.003; path *ab* = − 0.86, *SE* = 0.42; path *c* = − 1.07, *SE* = 0.45, *p* = 0.02; path *c*′ = − 1.93, *SE* = 0.54, *p* < 0.001, Figure 4h). This confirms that it is the representation of a hippocampal cognitive map that is critical for guiding spatial generalization and inference during the choice task and and the inference test. Furthermore, despite the fact that the spatial and the temporal kernel were correlated in most participants (average Pearson’s *r* = 0.58 ± 0.12), the neural effect as well as the degree to which behavior was influenced by either component could not be explained by a correlation between spatial and temporal kernels (Supplementary Figure S6).

### The representation of cognitive maps adapts to the task demands

In the choice task, rewards associated with the monsters were determined by their location in space and participants who had a better neural representation of the spatial map performed better in the inference tasks (Figure 4e). Yet, we also found evidence for a lingering effect of experienced temporal distances on choice.

We hypothesized that individuals adjust the degree to which they rely on one over the other map for guiding choice depending on the observed outcome contingencies. Indeed, a logistic function fitted to how individual weights changed over trials showed that in most participants the temporal component explained generalization behavior in the choice task better initially, but as the choice task progressed, spatial knowledge became more influential (Figure 5a). The slope of this logistic function was particularly steep for participants who performed better in the choice task (Figure 5a) as well as in the inference test (Figure 5b, *r* = −0.44, *p* = 0.002).

**Figure 5.**
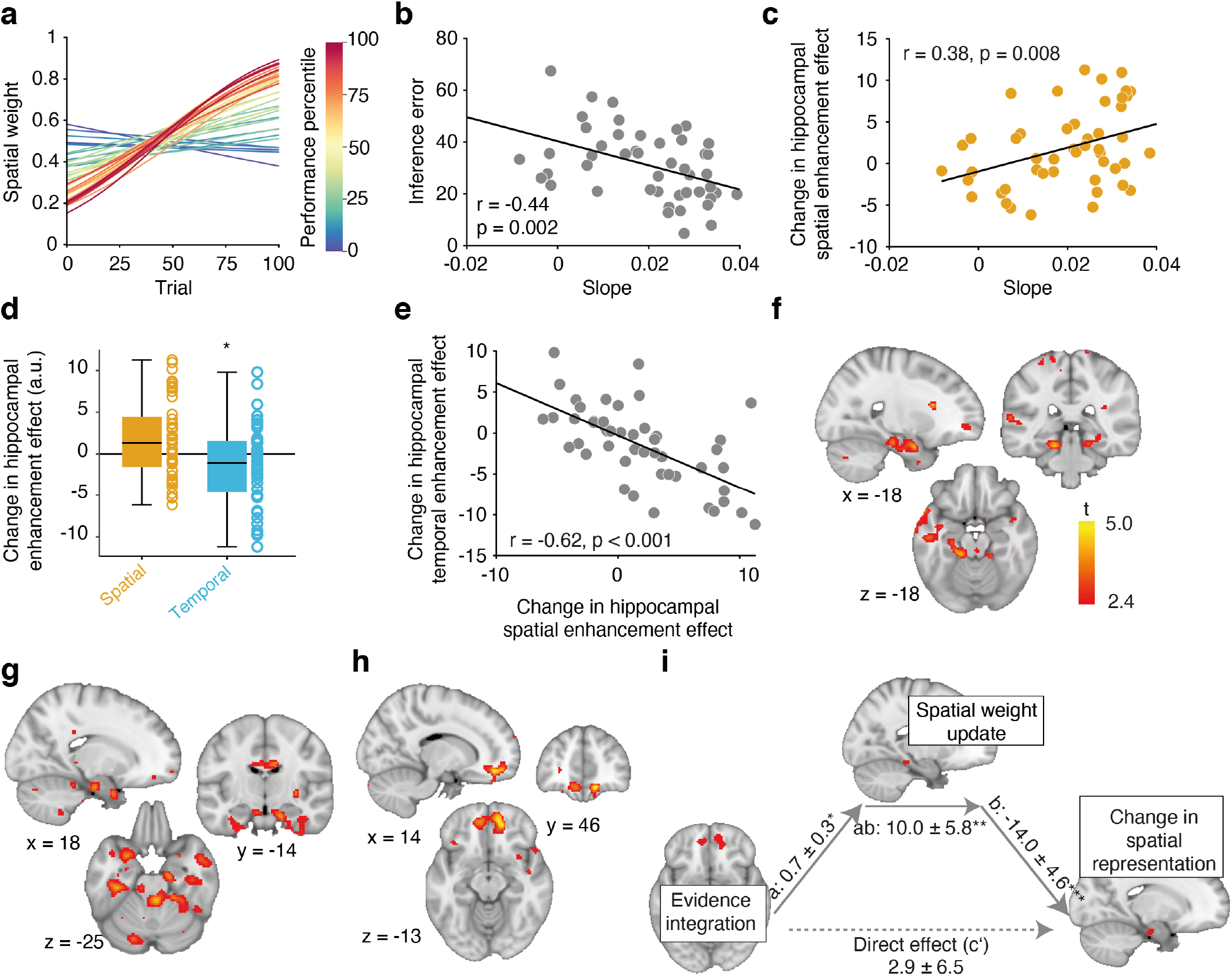
Hippocampal cognitive maps adapted to the task demands. **a** Logistic functions for each participant fitted to how individual spatial weights changed over trials. Curves are colored according to a participant’s relative performance in the choice task. **b** Correlation between the slopes of the estimated logistic function depicted in **a** and the inference error. **c** Correlation between the slopes of the logistic function and the change in the hippocampal spatial enhancement effect extracted from the ROI depicted in Figure 4a). **d** Change in spatial and temporal enhancement cross-stimulus enhancement effects in the ROI depicted in Figure 4a). Because the ROI was defined based on the existence of a spatial enhancement effect in session 3, the spatial effect is biased and displayed for visualization only. **e** Correlation between the change in the hippocampal spatial and temporal enhancement effects. Both were extracted from the ROI depicted in Figure 4a). **f** Whole-brain analysis depicting the update in spatial weights at the time of feedback. **g** Whole-brain analysis depicting voxels where the increase in the spatial cross-stimulus enhancement effect across participants correlates with the size of the hippocampal spatial weight update during the choice task as shown in **f. h** Whole-brain analysis depicting voxels where the difference in unsigned prediction errors as computed based on the temporal versus the spatial map correlates with with the size of the hippocampal spatial weight update during the choice task as shown in **f. i** Mediation path diagram for the change in the hippocampal spatial cross-stimulus enhancement effect extracted from the ROI depicted in Figure 4a as predicted by the OFC evidence integration signal and the hippocampal spatial weight update. **f-h** are thresholded at *p* < 0.01, uncorrected for visualization. * *p* < 0.05; ** *p* < 0.01; *** *p* < 0.001

We reasoned that this might reflect changes in the representation of the neural map over the course of the choice task. If this is the case, then participants who showed a larger increase in the contribution of spatial knowledge on choices, i.e. a steeper slope in the logistic regression, should also show a larger increase in the neural representation of the spatial map from day 2 (before the choice task) to day 3 (after the choice task). To test this, we extracted parameter estimates from the same region of interest we used for the analyses in Figure 4 for session 2 (before the choice task) and session 3 (after the choice task) and correlated the difference with the slope of the logistic function. The positive relation we found suggests that participants whose behavior in the choice task was characterized by marked increases in the reliance on the spatial map during choice also showed a larger increase in the neural representation of the spatial map (Figure 5c, *r* = − 0.44, *p* = 0.002). In the same region, the temporal map decreased significantly across participants (Figure 5d, *t*(47) = − 2.1, *p* = 0.04) and the change in the spatial map representation was negatively correlated with the change in the temporal map representation (Figure 5e, *r* = − 0.62, *p* < 0.001), suggesting that in participants where the spatial map representation became stronger, the temporal map representation became weaker.

We reasoned that this change in representation might be driven by a neural signal reflecting the degree to which either map was task-relevant during the choice task. To test this hypothesis, we first set up a GLM which included a parametric regressor that reflected the difference in the degree to which the spatial map influenced choice from one trial to the next. This identified a region in the left hippocampus tracking the trial-by-trial change in the degree to which the spatial dimension guided choice (Figure 5f, *t*(47) = 4.14, *p* = 0.02, [−18, −32, −18]).

If this neural weight update signal led to an increase in the neural representation of the relevant map, then participants with stronger hippocampal weight updating signals should display a larger change in hippocampal representation of the spatial map from day 2 to day 3. To test where the spatial weight updating signal correlated with a change in the spatial map representation, we looked for changes in the spatial map representation from session 2 to session 3 across the whole brain, and included the parameter estimates extracted from the hippocampal ROI reflecting the spatial weight update as a covariate. This analysis revealed a significant positive effect in the left hippocampal formation (Figure 5g, *p* = 0.018, *t*(47) = 4.21, [18, − 14, − 25]), suggesting that participants whose hippocampus tracked the spatial weight updates during the choice task also updated the representation of the spatial map in the hippocampus.

The changes in the composition of the hippocampal map likely reflect a representation learning process that was driven by the experienced reward contingencies in the choice task. To test whether any brain region tracks the evidence that the observed outcomes were generated by either of the two maps, we calculated the trial-wise unsigned prediction errors for each outcome separately for the spatial and the temporal map. The difference between these two prediction errors indicates how much more expected an outcome was according to the spatial as compared to the temporal map. We then set up a GLM that modeled this difference between spatial and temporal prediction errors at feedback time. Areas reflecting the evidence for the spatial over the temporal map should respond positively on trials where the spatial map made more accurate predictions than the temporal maps. We reasoned that, if there is a relationship between this signal and the spatial updating signal, then participants whose hippocampal weight updating signal was stronger should also show more of such an evidence tracking signal, and therefore included the parameter estimate extracted from the hippocampal ROI as a covariate. The only region where an evidence integration signal covaried with the hippocampal updating signal was the medial orbitofrontal cortex (Figure 5h, *p* = 0.03, [14, 46, − 13], family-wise error corrected on the cluster level).

In line with the observation that the OFC adapts behavior by changing associative representations in other brain regions^44^, the orbitofrontal evidence signal may thus align task representation with observed outcomes. By signalling the degree to which either map is task-relevant, spatial weights may be updated during the choice task, which in turn leads to an update of the spatial map representation itself. To test this assumption, we investigated whether the spatial weight update in the hippocampus formally mediated the relationship between the evidence integration signal in the OFC and the hippocampal changes in the spatial map representation. The fact that the OFC signal and the hippocampal spatial weight update was significant (path *a* = 0.7, *SE* = 0.3, *p* = 0.02) is not surprising, since the ROI was identified based on voxels where the corresponding covariate explains some variance. However, the effect of the spatial weight updating signal on the change in representation remains significant if we control for the OFC signal (path *b* = 14.0, *SE* = 4.6, *p* = 0.0003). Furthermore, there is a relationship between the OFC signal and the change in hippocampal map representation (path *c* = 13.1, *SE* = 6.6, *p* = 0.03), which can be fully accounted for by the hippocampal weight update (path *c*′ = 2.9, *SE* = 6.5, *p* = 0.6, path *ab* = 10.23, *SE* = 5.71, *p* = 0.007, Figure 4h). Hence, participants with the largest OFC evidence integration signal at feedback time exhibited the largest updates in spatial weights in the hippocampus, which in turn related to a larger change in the neural representation of the spatial map. This suggests a role for OFC signal in adjusting the use of an appropriate map to the current task demands, and an associated behavioral change.

## Discussion

The hippocampal formation is known to organize relationships between events in cognitive maps, thought to be critical for generalization and inference. However, the neural and computational mechanisms underlying the ability to use cognitive maps for generalization remains unknown in situations where stimuli are embedded in multiple relational structures. Here, we combined virtual reality, computational modeling and fMRI to demonstrate that the hippocampus extracts both spatial and temporal stimulus relationships from experience during navigation in a virtual arena. The strength of each neural representation was related to the degree to which it influenced behavior in an independent choice task. Notably, the OFC tracked the evidence that outcomes observed in the choice task were consistent with the predictions made by the spatial and the temporal cognitive map, which led to corresponding adjustments of the hippocampal map representation.

Participants learned to locate stimuli in a virtual arena. Because most individuals chose non-random behavioral policies for exploring the arena, stimulus relationships could be characterized both in terms of spatial distance as well as temporal co-occurrence. We found that the hippocampal formation extracted both types of relationships and represented those in clusters well-known to represent distances to goals^45^, goal direction signals^46^ as well as associative distances between objects forming a non-spatial graph^23^. Notably, the degree to which either map was represented in this region determined the degree to which participant’s generalization behavior in a later choice task was influenced by the corresponding map. This demonstrates a clear link between hippocampal map representations and their use for guiding generalization in decision making. It also shows that this system efficiently deals with higher-dimensional relational structures and can combine information from multiple maps for guiding choice.

We found substantial inter-individual differences in terms of the degree to which participants represented the spatial and temporal relationships a stimulus was embedded in neurally, and were influenced by those dimensions during choice. Indeed, in participants whose choices were influenced by the spatial or the temporal map, we found a cross-stimulus enhancement effect for spatial or temporal stimulus relationships, respectively. In participants whose choices were not influenced by those dimensions, on the other hand, the opposite was true: responses to a stimulus were suppressed if the preceding stimulus was close in space or time. Often, repetition suppression effects are more common than repetition enhancement effects in fMRI adaptation paradigms^40^. However, behavioral relevance can influence the directionality of an fMRI adaptation effect. For example, while repetition suppression effects are typically observed in the hippocampus when a stimulus that is irrelevant for the task at hand is repeated, repetition enhancement effects can be observed in the same region when a stimulus is task-relevant^47^. It is therefore conceivable that what a participant considered the relevant stimulus dimension was enhanced, while the irrelevant dimension was suppressed. In the context of our experiment, it was more adaptive to generalize along spatial rather than temporal distances, since spatial distances were used for creating reward contingencies in the first place. The more a participant therefore succeeded in enhancing the spatial dimension and suppressing the temporal dimension, the better they performed in the task.

Furthermore, participant choices became increasingly more influenced by spatial relational knowledge as the choice task progressed, suggesting that which map is used for guiding choice can be adaptively adjusted to the current task demands. This effect was driven by an OFC evidence integration signal, indexing the difference in accuracy of the predictions made by a spatial compared to a temporal model at feedback time. Participants whose OFC responded more strongly also showed a larger spatial weight updating signal in the hippocampus at feedback, which was in turn related to a stronger increase in the representation of the spatial map from before to after the choice task. This suggests that the OFC tracks the evidence that the currently observable state of the world was driven by either of the two maps, and updates the degree to which either influences behavior accordingly.

Our findings are consistent with the proposed function of the OFC to represent state spaces, in particular in situations where the current state of the world is not readily observable and must be inferred^48^. The OFC is also typically involved in situations where participants need to adjust their behaviour when outcome contingencies change^30^ or when memory responses require an arbitration between hippocampal and striatal inputs^49^. For example, reversal learning or outcome devaluation, where previously acquired cue–outcome and response–outcome associations need to be adapted, rely on an intact OFC^50^.

Importantly, our results also shed light on the interaction between OFC and the hippocampus. In line with previous observations indicating a relation between state representations in OFC and the hippocampus^31, 51, 52^, our results indicate that OFC might play an active role in learning state presentations in the hippocampus through experience^53^. Future experiments should assess whether similar adjustments can also be observed when temporal rather than spatial stimulus relationships govern the reward distribution, or when rewards are governed by a compositional, spatio-temporal map.

In conclusion, our results suggest that the hippocampus represents different dimensions of experienced relationships between stimuli such as space and time in parallel cognitive maps. The degree to which each one is used for guiding choice is governed by an OFC evidence integration signal. The OFC drives a spatial updating signal in the hippocampus, which is in turn related to a change in the representation of the spatial map. This provides a mechanistic insight into the way in which appropriate stimulus dimensions are selected for guiding decision making in multi-dimensional environments.

## Online Methods

### Participants

52 neurologically and psychiatrically healthy participants took part in this study (mean age 26.8 ± 3.8 years, 20-34 years old, 27 male). Participants were recruited using the participant database of the Max Planck Institute for Human Cognitive and Brain Sciences. Due to a scanner defect, three participants could not complete the last day. One participant was excluded due to problems during the preprocessing. 48 participants therefore entered the analyses. Two of those participants did not do the arena task at the end of the experiment, but their data was included in all other analyses. The study was approved by the ethics committee at the Medical Faculty at the University of Leipzig (221/18-ek) and all participants gave written informed consent prior to participation.

### Experimental procedure

The experiment consisted of three parts performed on three subsequent days. On day 1, participants learned the stimulus distribution in a virtual arena. On day 2, we assessed the stimulus representation in the fMRI scanner. On day 3, participants performed a choice task to learn the rewards associated with each stimulus in the scanner. Afterwards, we again assessed the stimulus representations in the scanner. The sessions are described in more detail below. The exploration and object location memory task were coded using the virtual reality software package Vizard (Version 4, Santa Barbara, CA: WorldViz LLC). All other tasks were written in custom-written Matlab scripts using Psychtoolbox. Imaging data was preprocessed using fmriprep. Imaging and behavioural analyses were carried out with Matlab.

#### Day 1

Participants were first familiarized with the stimuli by being presented with the monsters one-by-one on the screen. They could click through the stimuli to proceed to the next one. Participants were then instructed that they would be asked to learn where each monster belongs in space, and that this knowledge would be important for collecting points in later sessions. Monsters were distributed in a circular arena with a virtual radius of 15m (Figure 1A). Which monster was presented in which location was randomized across participants. 5 distinct trees were located behind the wall surrounding the arena, which functioned as landmarks. The location of the trees was randomized in such a way that one tree occurred at a random position in every 72deg block in each participant. Tree locations were fixed across all experimental session.

Participants then learned the location of stimuli in space by navigating around a virtual arena (Figure 1E) in multiple blocks. Each block consisted of an exploration phase and an object location memory task. In the exploration phase, participants navigated around the arena in any way they liked and for as long as they wanted. Whenever a participant approached a monster (i.e. they entered a 3 m radius around the monster location), it became visible and slowly turned around its own axis. This means that participants never saw all monsters at the same time. After each exploration phase, participants performed an object location memory task. In this task, participants were cued with a monster and had to navigate to the corresponding location (Figure 1F). Feedback indicated how close to the correct location a monster was positioned (<3m, <5m, <7m, <9m, >9m). In each block, every monster had to be positioned once. The order was randomized. If performance reached a pre-specified performance criterion of <3m drop error averaged across all monsters (corresponding to <10% error) in a block, the session terminated if a participant had completed at least five blocks. Participants performed a minimum number of 5 and a maximum number of ten blocks of this task to ensure that they had a good knowledge of the stimulus distribution.

#### Day 2

Before the scanning session, participants had another opportunity to explore the monster locations freely, followed by one more round of the object location memory task with feedback.

Subsequently, we assessed the monster representations in the scanner using a picture viewing task. Here, participants were presented in the fMRI scanner with the monsters in a random order on a red or a blue background. Participants were instructed to view the images attentively. Occasionally (once after each monster on each background color), two monsters were presented simultaneously and participants had to indicate which of the two monsters was located closer in space to the monster they had seen immediately before the two monsters. Participants received no feedback. The purpose of this task was to ensure that participants would always evoke the location a monster was embedded in during the stimulus presentations. Correct answers were rewarded with 0.10 EUR. Participants were instructed that the background color was irrelevant for performing the task. Each monster was presented 6 times on each background color (red, blue) per block, resulting in 144 stimulus presentations in each block. Participants completed three blocks of this task. Stimulus sequences were generated pseudo-randomly using a genetic algorithm with the following constraints: Each stimulus in each context occurred the same number of times per block and no monster-monster transition was presented more than once.

After the scanning session, another round of the object location memory task was performed without feedback to assess participants’ memory for the monster locations.

#### Day 3

Before the scanning session, another round of the object location memory task was performed without feedback to assess participants’ memory for the monster locations.

In the scanner, participants then performed a choice task. Here, they were presented with pairs of monsters and instructed to select the monster that would lead to the highest reward. The reward distribution was related to the position of the monsters in space and the context as indicated by the background color (Figure 1H). Participants were instructed that they would receive similar amounts of points for monsters located near each other in space. They learned the two value distributions in a blocked fashion, with ten trials of choices in context 1 alternating with ten trials of choices in context 2. Background colors and contexts were counterbalanced across participants. Value distributions were selected such that pairwise spatial distances and pairwise value differences across both contexts were not significantly correlated and that the overall value across all objects was similar across the two contexts.

Two objects in each context (‘inference objects’) could never be chosen during the choice task (Figure 1B). These were later used to assess whether participants were able to combine information about rewards with information about the relationship between monsters to infer stimulus values that were never directly experienced. Critically, the value of one inference object per context was high (71 and 72) and the value of the other inference objects was low (3 and 13).

After the choice task, three blocks of the picture viewing task were performed in the scanner. This time, the background colour indicated the relevant context and participants were instructed to think about each monster’s location in space and its associated value. Occasionally (once after each monster on each background color), two monsters were presented simultaneously and participants had to indicate which of the two monsters was located closer in space to the monster they had seen immediately before the two monsters or which monster had a more similar value. Which task was to be performed was indicated with a symbol presented above the two options. Correct answers were rewarded with 0.10 EUR. Stimulus sequences were the same as on day 2.

After the scanning session, another round of the object location memory task was performed without feedback to assess participants memory for the monster locations. This was followed by four brief tasks. (1) Participants had to indicate on a sliding scale from 0 to 100 how many points they would receive for each monster in each context, (2) Participants rated on a scale from “not at all” to “very much” how much they liked each monster, (3) Participants arranged monsters in an arena according to their similarity (Arena task 1), and (4) according to their spatial location (Arena task 2). In each task, the order in which monsters were presented was randomized across participants.

### Reimbursement

Participants were paid a baseline fee of 9 €/hour for the behavioral parts of the experiment and 10 €/hour for the fMRI sessions. In addition, participants could earn a monetary bonus depending on performance. Points accumulated during the choice blocks were converted into money (100 points = 0.1 €). Furthermore, each correct choice during the monster presentation block was rewarded with 0.10 €.

### Behavioral analysis

#### Object positioning task

The replacement error in the object location memory task was defined as the Euclidean distance between the drop location and the true object location. It was reported relative to the arena diameter.

#### Choice task

A correct choice was the choice corresponding to the object with the higher value.

#### Inference task

The inference error was defined as the root-mean-square error between the true inference values and the error ratings provided by a participant at the end of the study.

#### Arena task

The map reproduction error was defined as the root-mean-square error between the true z-scored spatial distances between the monsters in the virtual arena and the z-scored distances between the monster positions in the arena task. We z-scored the distances to ensure that they had a comparable range.

### Modeling

We used Gaussian process regression to model reward learning and generalization in the choice task. Gaussian processes (GPs) define probability distributions over functions *f* ∼ 𝒩 (*m*(**x**), *k*(**x, x**′)), where *m*(**x**) is the mean function, giving the expected function values **ŷ** at input points **x**, and *k*(**x, x**′) the covariance function, or kernel, defining how similar any pair of input points, **x** and **x**′, are. GPs can be updated to posterior distributions over functions by conditioning on a set of observed function outputs **y**. Here the posterior mean function is given by

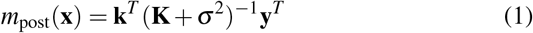

where **k** is the kernel matrix containing the covariance between training points and the evaluation points, **K** is the kernel matrix containing the covariance between all training points, and *σ*^2^ is a diagonal variance matrix.

The hypothesis that generalization is guided by a spatial cognitive map corresponds to equipping a GP model with a Gaussian (or Radial Basis Function) kernel, representing similarity as an exponentially decaying function of squared Euclidean distance. The Gaussian kernel defines similarity as follows:

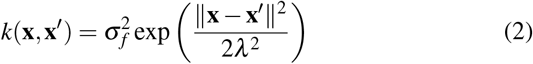

where 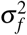 is a parameter controlling the degree to which the predictions differ from the mean, and *λ* is the lengthscale parameter, controlling how strongly input point similarity decays with distance. We obtained estimates of stimuli locations for every participant by performing path integration on their navigation runs.

To construct a kernel that corresponds to the hypothesis that temporal relations guided generalization, we started by computing a successor matrix **M** for every participant^33^. Each entry in the successor matrix **M**(*s, s*′) (Equation 3) contains the expected discounted number of future visits of stimulus *s*′, starting from a visit to stimulus *s*

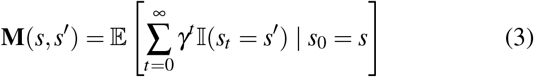

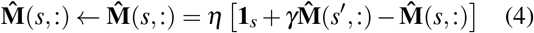

where *γ* is the discount factor and 𝕀 is the indicator function. The successor matrix can be approximated from a participant’s stimulus visitation history using a simple temporal-difference updating rule^54^ (Equation 4), where 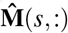 is the row corresponding to stimulus *s*, **1**_*s*_ is a vector of zeros except for the *s*th component which is a 1, and *η* is the learning rate. From **M** we computed the transition matrix **T** using the following equation (see Supplementary Note section for derivation):

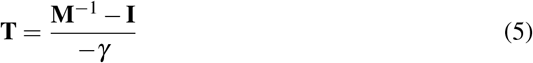

where **I** is the identity matrix. We enforced that **T** was symmetric by taking the pairwise maximum of the entries of its upper and lower triangles. From **T**, which describes the relevant participant’s probabilities of walking directly from one stimulus to another, we computed the diffusion kernel^34^ **K**, embodying the hypothesis that temporal relations guide generalizations (Equation 6).

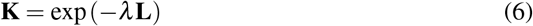

Here exp is matrix exponentiation, **L** is the normalized graph Laplacian which equals **I**−**T**, and *λ* is a lengthscale parameter analogous to that of the Gaussian kernel (Equation 2). To obtain the compositional kernel we took the average of the Gaussian and the diffusion kernel^55^, and to implement the mean tracker we used a GP model whose kernel was the identity matrix **I**.

To obtain the various GP models’ estimates of stimuli’s rewards at any given trial in the choice task, we conditioned them on all previously observed stimuli’s rewards for the relevant context up to that point, and computed the posterior mean using Equation 1. The differences in estimated rewards were used as single predictors of participant choices in a logistic mixed-effects model with a participant-specific random slope^38^, implemented in R using the `lme4`^56^ package. We optimized hyper-parameters to minimize the log-likelihood of producing the choice data using a grid-search.

For the Gaussian kernel, we optimized the lengthscale *λ*, for the diffusion kernel we optimized the learning rate *η*, and set the discount rate parameter *γ* to 0.9 and the lengthscale *λ* to 1. For the compositional, spatio-temporal kernel, we optimized both the Gaussian kernel’s lengthscale and the learning rate. The variance in Equation 1 was set 0.01 to improve numerical stability for matrix inversion. Using the best-fitting hyperparameter configurations, we performed a leave-one-out cross-validation (LOO-CV) procedure and obtained each model’s cross-validated log-likelihood of producing every choice in the data set. We then computed the posterior model frequencies and exceedance probabilities^57^, reported in Figure 3B.

We used the same procedure for modelling participants’ value judgements. Here, we made the GP models predict the values of all stimuli, based on all reward-observations the participants had made, respectively. The GPs were equipped with the best-fitting hyper-parameters (see Supplementary Note section) from the choice task. We then sought to predict participants’ value judgements for the different stimuli using the various value estimates as single predictors (plus an intercept) in separate linear mixed-effects models with a participant-specific random slope. We split the value judgements into two sets: One containing the value judgements of the inference objects, and another containing the value judgements of all monsters except the inference objects. Again, we performed LOO-CV to obtain model-specific log-likelihoods for all value judgements in the two data sets. Since the mean tracker could not generate predictions for the inference object any different from its prior mean function (which was 0), we used the average of the mean tracker’s value predictions for the non-inference objects as a baseline model. From the cross-validated log-likelihoods we computed the corresponding sets of model frequencies and exceedance probabilities.

To compute the effects of the spatial and temporal components on each participant’s choice behaviour, we fitted mixed-effects logistic regression models like the ones described above, using the estimated value differences generated by the spatial and temporal maps as individual predictors (using their respective best-fitting hyper-parameters) in the same model. Since the two predictors were correlated, we created two such models, one where the spatial value difference was the main predictor, and the second predictor was the temporal *minus* the spatial predictor, and a second model where this relation was inverted^58^. We aggregated the unsigned mixed effects (random effects plus the fixed effects) across these two models for all participants, which left us with the effects for the two maps. To compute the spatial weights, we calculated how big the spatial effects were in proportion to the total effects (spatial + temporal effects). The temporal weights were consequently 1 minus the spatial weights. To compute the slopes, we first obtained a weight for the spatial map for all trials, and for all participants. We computed these weights by estimating two models similar to the ones used to estimate participant-specific effects, this time including an interaction term with trial number as well. To obtain trial-specific spatial weights for all participants, we estimated how likely the spatial × trial interaction predictor was at predicting each individual choice compared to the temporal × trial interaction predictor, aggregating over our two models. We then fitted logistic slopes to each participant’s spatial weight time series, predicting single participants’ spatial weights from trial number, using logistic regression.

### fMRI data acquisition and pre-processing

Visual stimuli were projected onto a screen via a computer monitor. Participants indicated their choice using an MRI-compatible button box.

MRI data were acquired using a 32-channel head coil on a 3 Tesla Siemens Magnetom SkyraFit system (Siemens, Erlangen, Germany). fMRI scans were acquired in axial orientation using T2*-weighted gradient-echo echo planar imaging (GE-EPI) with multiband acceleration, sensitive to blood oxygen level-dependent (BOLD) contrast^59, 60^. Echo-planar imaging (EPI) with sampling after multiband excitation achieves temporal resolution in the sub-second regime whilst maintaining a good slice coverage and spatial resolution^59, 60^. We collected 60 transverse slices of 2-mm thickness with an in-plane resolution of 2 × 2 mm, a multiband acceleration factor of 3, a repetition time of 2 s, and an echo time of 23.6 ms. Slices were tilted by 90 deg relative to the rostro-caudal axis. The first five volumes of each block were discarded to allow for scanner equilibration. Furthermore, a T1-weighted anatomical scan with 1 × 1 × 1 mm resolution was acquired. In addition, a whole-brain field map with dual echo-time images (TE1 = 5.92 ms, TE2 = 8.38 ms, resolution 2 × 2 × 2.26 mm) was obtained in order to measure and later correct for geometric distortions due to susceptibility-induced field inhomogeneities.

#### Anatomical data preprocessing

Results included in this manuscript come from preprocessing performed using *fMRIPrep* 1.4.0^61, 62^ (RRID:SCR_016216), which is based on *Nipype* 1.2.0^63, 64^ (RRID:SCR_002502).

A total of 2 T1-weighted (T1w) images were found within the input BIDS dataset. All of them were corrected for intensity non-uniformity (INU) with `N4BiasFieldCorrection`^65^, distributed with ANTs 2.2.0^66^. The T1w-reference was then skull-stripped with a *Nipype* implementation of the `antsBrainExtraction.sh` workflow (from ANTs), using OASIS30ANTs as target template. Brain tissue segmentation of cerebrospinal fluid (CSF), white-matter (WM) and gray-matter (GM) was performed on the brain-extracted T1w using `fast`^67^. A T1w-reference map was computed after registration of 2 T1w images (after INU-correction) using `mri_robust_template`^68^.

Brain surfaces were reconstructed using `recon-all`^69^, and the brain mask estimated previously was refined with a custom variation of the method to reconcile ANTs-derived and FreeSurfer-derived segmentations of the cortical gray-matter of Mindboggle^70^. Volume-based spatial normalization to one standard space (MNI152NLin6Asym) was performed through nonlinear registration with `antsRegistration` (ANTs 2.2.0), using brain-extracted versions of both T1w reference and the T1w template. The following template was selected for spatial normalization: *FSL’s MNI ICBM 152 non-linear 6th Generation Asymmetric Average Brain Stereotaxic Registration Model*^71^ [RRID:SCR_002823; TemplateFlow ID: MNI152NLin6Asym].

#### Functional data preprocessing

For each of the 7 BOLD runs per subject (across all tasks and sessions), the following preprocessing was performed. First, a reference volume and its skull-stripped version were generated using a custom methodology of *fMRIPrep*. A deformation field to correct for susceptibility distortions was estimated based on a field map that was co-registered to the BOLD reference, using a custom workflow of *fMRIPrep* derived from D. Greve’s `epidewarp.fsl` script and further improvements of HCP Pipelines^72^. Based on the estimated susceptibility distortion, an unwarped BOLD reference was calculated for a more accurate co-registration with the anatomical reference. The BOLD reference was then co-registered to the T1w reference using `bbregister` (FreeSurfer) which implements boundary-based registration^73^. Co-registration was configured with nine degrees of freedom to account for distortions remaining in the BOLD reference. Head-motion parameters with respect to the BOLD reference (transformation matrices, and six corresponding rotation and translation parameters) are estimated before any spatiotemporal filtering using `mcflirt`^74^.

BOLD runs were slice-time corrected using `3dTshift` from AFNI 20190105^75^. The BOLD time-series (including slice-timing correction when applied) were resampled onto their original, native space by applying a single, composite transform to correct for head-motion and susceptibility distortions. These resampled BOLD time-series will be referred to as *preprocessed BOLD in original space*, or just *preprocessed BOLD*. The BOLD time-series were resampled into standard space, generating a *preprocessed BOLD run in [‘MNI152NLin6Asym’] space*. First, a reference volume and its skull-stripped version were generated using a custom methodology of *fMRIPrep*.

Additionally, several confounding time-series were calculated based on the *preprocessed BOLD*: framewise displacement (FD), DVARS and three region-wise global signals. FD and DVARS are calculated for each functional run, both using their implementations in *Nipype*^76^. The three global signals are extracted within the CSF, the WM, and the whole-brain masks. Additionally, a set of physiological regressors were extracted to allow for component-based noise correction *CompCor*^77^. Principal components are estimated after high-pass filtering the *preprocessed BOLD* time-series (using a discrete cosine filter with 128s cut-off) for the two *CompCor* variants: temporal (tCompCor) and anatomical (aCompCor). tCompCor components are then calculated from the top 5% variable voxels within a mask covering the subcortical regions. This subcortical mask is obtained by heavily eroding the brain mask, which ensures it does not include cortical GM regions. For aCompCor, components are calculated within the intersection of the aforementioned mask and the union of CSF and WM masks calculated in T1w space, after their projection to the native space of each functional run (using the inverse BOLD-to-T1w transformation). Components are also calculated separately within the WM and CSF masks. For each CompCor decomposition, the *k* components with the largest singular values are retained, such that the retained components’ time series are sufficient to explain 50 percent of variance across the nuisance mask (CSF, WM, combined, or temporal). The remaining components are dropped from consideration. The head-motion estimates calculated in the correction step were also placed within the corresponding confounds file. The confound time series derived from head motion estimates and global signals were expanded with the inclusion of temporal derivatives and quadratic terms for each^78^.

Frames that exceeded a threshold of 0.5 mm FD or 1.5 standardised DVARS were annotated as motion outliers. All resamplings can be performed with *a single interpolation step* by composing all the pertinent transformations (i.e. head-motion transform matrices, susceptibility distortion correction when available, and co-registrations to anatomical and output spaces). Gridded (volumetric) resamplings were performed using `antsApplyTransforms` (ANTs), configured with Lanczos interpolation to minimize the smoothing effects of other kernels^79^. Non-gridded (surface) resamplings were performed using `mri_vol2surf` (FreeSurfer).

### fMRI data analysis

We implemented three types of event-related general linear models (GLMs) in SPM 12 to analyze the fMRI data. All GLMs included a button press regressor as a regressor of no interest. All regressors were convolved with a canonical haemodynamic response function. Because of the sensitivity of the blood oxygen level-dependent signal to motion and physiological noise, all GLMs included frame-wise displacement, six rigid-body motion parameters (three translations and three rotation), six anatomical component-based noise correction components (aCompCorr) and four cosine regressors estimated by fmriprep as confound regressors for denoising. Each block was modeled separately within the GLMs.

The first GLM contained separate onset regressors for each of the twelve objects. By modeling each object separately, we could account for any object-specific differences in activity driving the main effects and focus on distance-dependent modulations that ride on top of those object-specific differences in activation. Each onset regressor was accompanied by two parametric regressors. These corresponded to the distance to the object presented immediately before the current object according to the spatial kernel and distance to the immediately preceding object according to the temporal kernel. Both parametric regressors were zscored, but not orthogonalized, so that any shared variance would be discarded. Trials where the same object was repeated were modeled separately and objects immediately following a choice were excluded. Furthermore, the GLM contained an onset regressor for the choice trials. This was accompanied by two parametric regressors, reflecting chosen and an unchosen distance between the two objects and the preceding object. Each of the three blocks were modeled separately.

The second and third GLM modeled events during the choice task. Here, three onset regressors were included, one indicating the choice period, the second one indicating feedback times and the third one corresponding to button presses. The duration of each event corresponded to the actual duration during the experiment. The choice period regressor was accompanied by two parametric modulators reflecting chosen and unchosen values of the objects as estimated by the winning model. Both were demeaned, but not orthogonalized.

In the second GLM instead, the feedback regressor was accompanied by a spatial weight updating signal. A trial-by-trial estimate of the influence of the spatial map on the choices was estimated, and the demeaned trial-by-trial difference was included as a parametric modulator.

In the third GLM, the feedback regressor was accompanied by a parametric regressor reflecting a prediction error difference signal. The reward prediction error was estimated separately for the spatial and the temporal map, and the demeaned difference between the absolute prediction errors was included as a parametric regressor.

The contrast images of all participants from the first level were analysed as a second-level random effects analysis. We report all our results in the hippocampal formation, as this was our a priori ROI, at an uncorrected cluster-defining threshold of *p* < 0.001, combined with peak-level family-wise error (FWE) small-volume correction at *p* < 0.05. For the SVC procedure, we used a mask comprising hippocampus, entorhinal cortex and subiculum (Supplementary Figure S5). Activations in other brain regions were only considered significant at a level of *p* < 0.001 uncorrected if they survived whole-brain FWE correction at the cluster level (*p* < 0.05). Results in the orbitofrontal cortex in 5h are reported at a cluster-defining threshold of *p* < 0.01 uncorrected, combined with a whole-brain FWE-corrected significance at the cluster level of *p* < 0.05. While we used masks to correct for multiple comparisons in our ROI, all statistical parametric maps presented in the manuscript are unmasked and thresholded at *p* < 0.01 for visualization.

To relate neural effects to behavioral parameters and to each other, we defined the following ROIs: spatial hippocampal map in session 3 from GLM 1, Figure 4a; hippocampal spatial weight update from GLM 2, Figure 5f; change in hippocampal map representation from session 2 to session 3 with hippocampal spatial weight update as covariate from GLM 1, Figure 5g; and OFC evidence integration signal with hippocampal spatial weight update as covariate from GLM 3 5h. All voxels exceeding a threshold of *p* < 0.001 were included in an ROI if the cluster survived correction for multiple comparisons.

To estimate how much an effect co-varied with behavioral effects, we included spatial and temporal weights, respectively (Figure 4f), as well as the inference error (Figure 4g) as a covariate on the second level and tested for significant effects. In Figure 5g and h, we included the parameter estimate reflecting the size of the hippocampal spatial weight update signal (Figure 5f) as a covariate.

### Mediation analysis

We used the Mediation and Moderation Toolbox^42, 43^ to perform two single-level mediation analyses (Figures 4h and 5i). The total effect of the independent variable X on the dependent variable Y is referred to as path c. That effect is then partitioned into a combination of a direct effect of X on Y (path c’), and an indirect effect of X on Y that is transmitted through a mediator M (path ab). We also estimated the relationship between X and M (path a) as well as between M and Y (path b). This last path “b” is controlled for X, such that paths “a” and “b” correspond to two separable processes contributing to Y. We determined two-tailed uncorrected p values from the bootstrap confidence intervals for the path coefficients^43^.

To test whether the spatial weights mediate the effect of hippocampal spatial map on the inference error, we defined X as each individual’s parameter estimate from the hippocampal ROI encoding the spatial map (ROI based on Figure 4a). The mediator M corresponded to each participant’s spatial weight as estimated by the model fit to the choice data. The outcome variable Y was defined as a participant’s inference error.

To test for a significant mediation linking the OFC evidence integration signals (X) to the change in hippocampal spatial map (Y), we extracted parameter estimates from an orbitofrontal ROI tracking the evidence that an outcome is predicted by either of the two maps (X, ROI based on Figure 5h) and related this to the change in spatial representation in the left hippocampus (Y, ROI based on Figure 5g) via the spatial updating signal in the right hippocampus (M, ROI based on Figure 5f).

## Data availability

Source data to reproduce the figures and unthresholded group-level statistical brain maps from neuroimaging analyses will be made openly available upon publication.

## Code availability

Task, analysis and computational modeling code will be made publicly available on github upon publication.

## Acknowledgements

We would like to thank Jacob Bellmund for helpful comments on the manuscript, Joshua Julian for providing example code for the VR experiment and Kerstin Träger and Nicolas Filler for help with data collection. We thank the University of Minnesota Center for Magnetic Resonance Research for the provision of the multiband EPI sequence software.

This work is supported by the Max Planck Society. TS and ES are supported via the Mini Graduate School on “Compositionality in Minds and Machines” from the Deutsche Forschungsgemeinschaft under Germany’s Excellence Strategy–EXC2064/1–390727645. ES is further supported by an Independent Max Planck Research Group grant awarded by the Max Planck Society. NWS is supported by an Independent Max Planck Research Group grant awarded by the Max Planck Society and a Starting Grant from the European Union (ERC-2019-StG REPLAY-852669). CFD is supported by the Max Planck Society, the European Research Council (ERC-CoG GEOCOG 724836), the Kavli Foundation, the Jebsen Foundation, the Centre of Excellence scheme of the Research Council of Norway – Centre for Neural Computation (223262/F50), The Egil and Pauline Braathen and Fred Kavli Centre for Cortical Microcircuits, and the National Infrastructure scheme of the Research Council of Norway – NORBRAIN (197467/F50).

## Author contributions statement

M.M.G., N.W.S. and C.F.D. conceived the experiment, M.M.G. developed the tasks and acquired the data, all authors planned the analyses, M.M.G. and T.S. analyzed the data, T.S. and E.S. performed the computational modeling, all authors discussed the results, M.M.G. and T.S. wrote the manuscript with input from all authors.

