## Supplemental Material for "Hippocampal spatio-temporal cognitive maps adaptively guide reward generalization"

### Supplementary Information

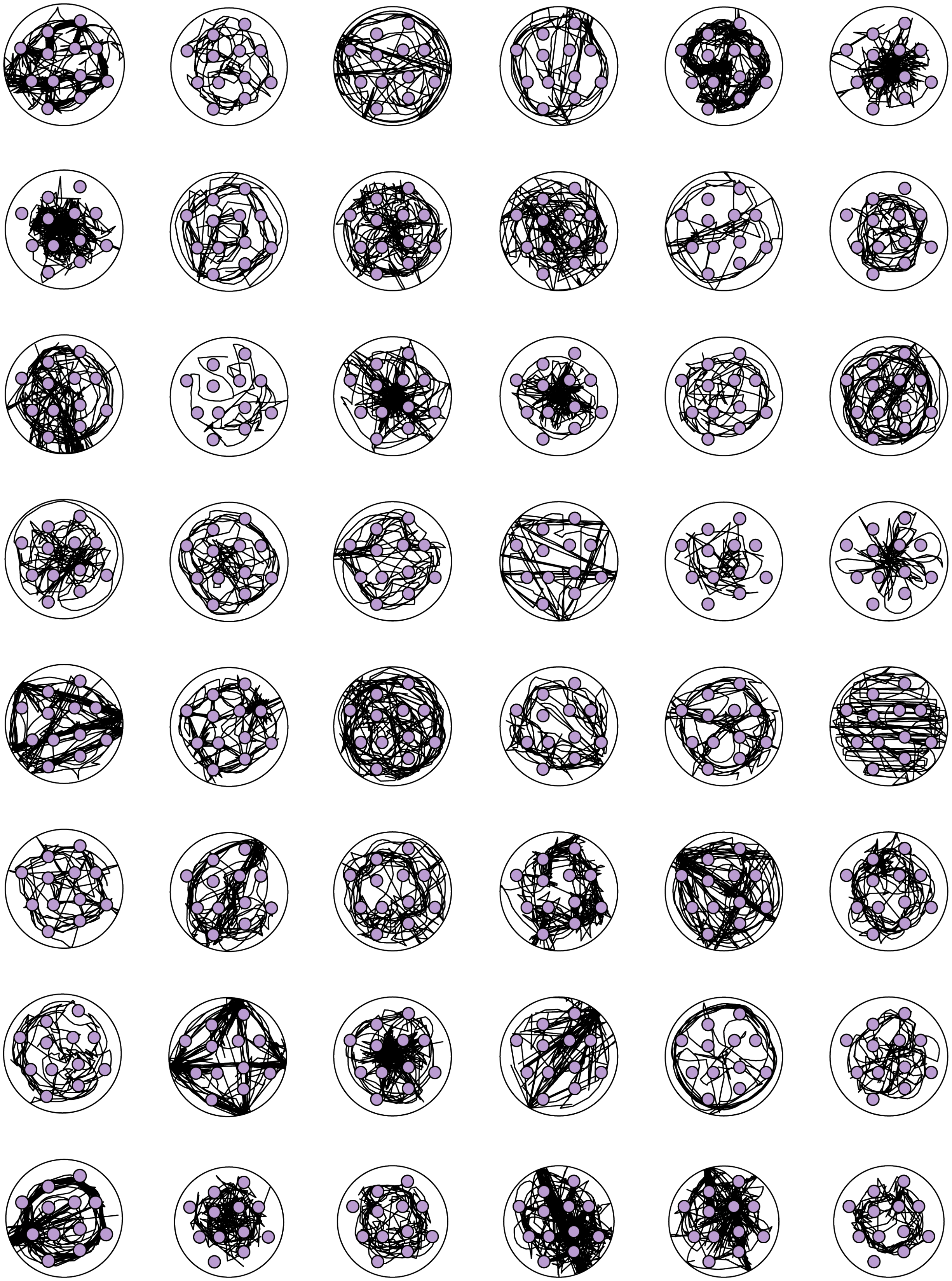

**Supplementary Figure S1. Exploration paths on day 1 in each individual.** Each panel represents the exploration trajectories concatenated across exploration blocks on day 1 in one participant. Purple indicates the stimulus locations and black the participant´s trajectory.

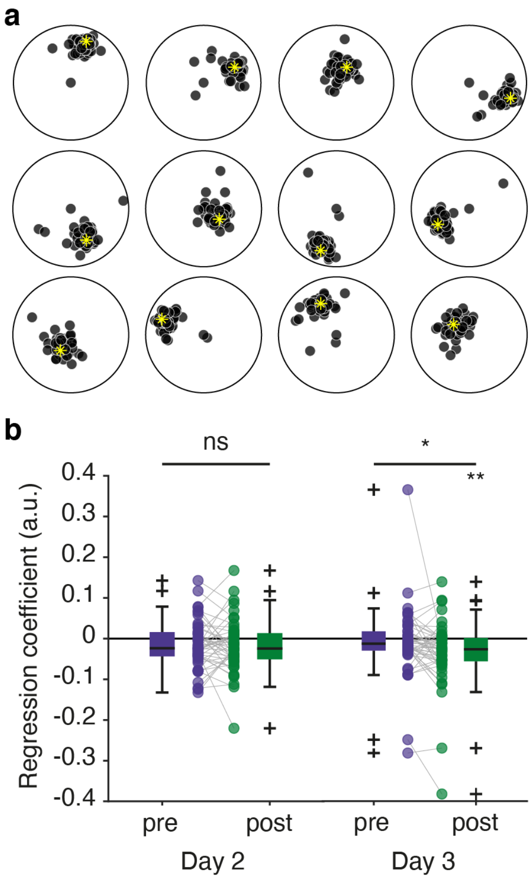


**Supplementary Figure S2. Object positioning after learning. a** Each panel displays the data for one object. Yellow indicates the true object position. Black indicates the drop location for each participant. The replacement error is defined as the Euclidean distance between the true location and the drop location. Visualized is the data from the last object location memory task block on day 1, i.e. at the end of learning. **b** Linear regression of values on replacement error. On day 2 as well as on day 3 before the choice task, there was no relationship between values participants learned to associate with each object and replacement error (all p values > 0.05). This is not surprising, since participants only learned the value associations on day 3. On day 3 after the choice task, the replacement error was smaller the higher the reported value of an object ($t(47) = -2.9, p = 0.005$). The difference between value-dependent performance pre and post choice was also significant on day 3 ($t(47) = 2.26, p = 0.03$), but not on day 2 ($t(47) = 0.27, p = 0.79$). This suggests that participants' memory expression was more accurate around valuable objects compared to less valuable ones after participants learned to associated objects with values. We used the average values that participants reported at the end of the study on day 3 as predictors. For inference objects, only the value experienced in the other context was considered.

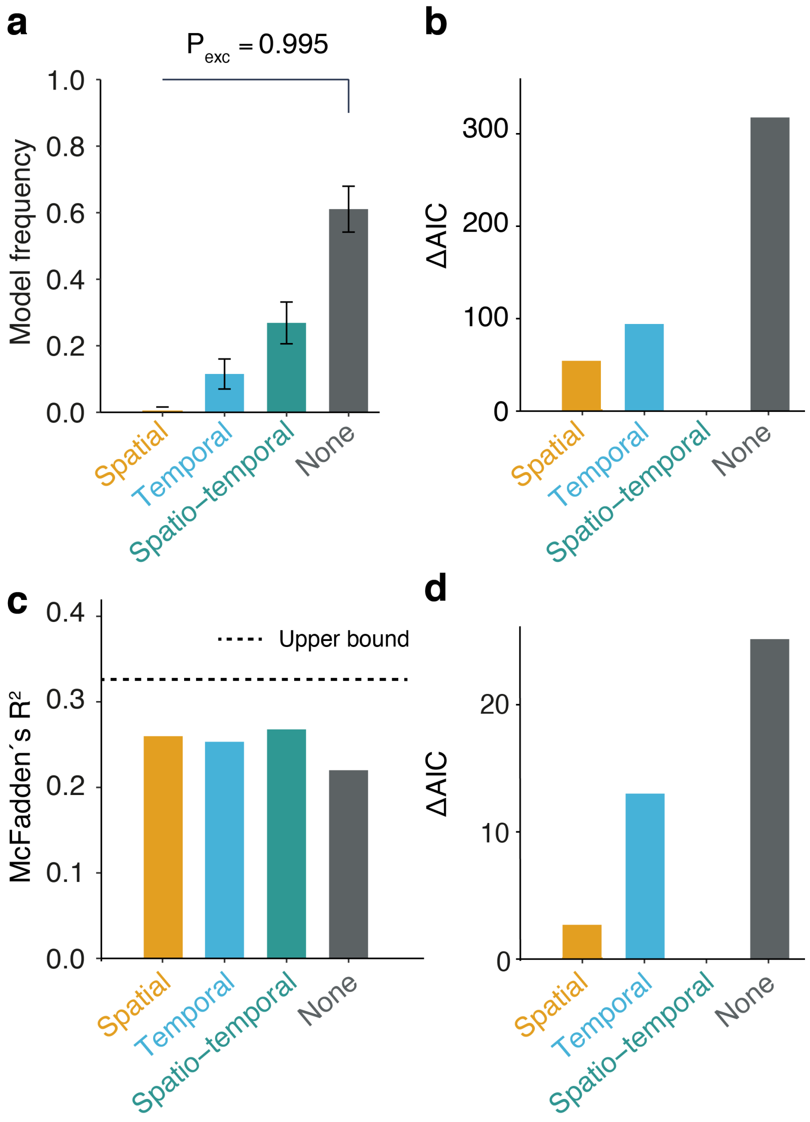


**Supplementary Figure S3. Full modeling results. a** Model frequencies for predicting participants' value ratings for the experienced objects at the end of the study. The winning model does not generalize about value. **b** Model AIC differences for the choice task. **c** Models' McFadden's $R^2$ for the choice task. This statistic quantifies how likely a model is to produce the data relative to a random model, where a score of 1 means that the model is infinitely more likely to produce the data, and a score of 0 means the model is as likely as the random model. The dashed line represents the score of a model that uses the true value difference between options as a predictor. This model was only tested on trials where participants had observed the value of both options, and whose score therefore approximates an upper bound on how accurately one can predict participant choices assuming one has access to their beliefs about value and perfect memory, relative to a chance levels. **d** Model AIC differences for predicting participants' value ratings for the inference objects.

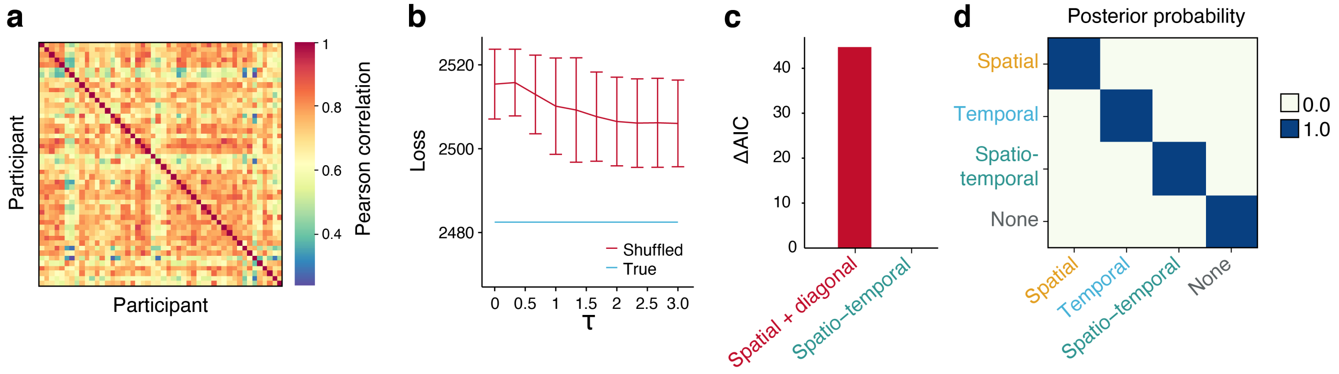


**Supplementary Figure S4. Model recovery. a** The pairwise correlation between all participants' temporal kernels estimated with a learning rate of 0.4125, which gave the best fit for the temporal model. We flattened each participant's $12 \times 12$ temporal kernel matrix into a vector and computed Pearson's correlation coefficient $r$ between all pairs of vectors. **b** Predicting reward generalization using participants' own temporal kernel yields substantially better fits to their choice behaviour (blue line) than predicting generalization using another randomly picked participant's temporal kernel (red line). Error bars are standard deviations of negative log-likelihood of 10 sampled random assignments. See Supplementary Note section  for procedure. **c** To verify that the predictive performance of the spatio-temporal model was not an artifact of the kernel composition procedure per se, we compared the spatio-temporal model against a model using a composition of a spatial kernel and the identity matrix. The spatio-temporal kernel produced a substantially better fit to participant choices, indicating that both the spatio-temporal model's components captures something important about how participants generalized. **d** We performed a model recovery analysis for our computational models, using their own best-fitting hyper-parameters. We first simulated choice behaviour from our models based on the choices that the participants encountered in the experiment. As such, we obtained 4800 simulated decisions from our models. The temporal and spatio-temporal models used the temporal kernel of the participant at the corresponding trial. Choices were made deterministically to maximize expected reward, where the expected reward was estimated from previous observations. After each choice, the models received a reward which they used to condition predictions about rewards for subsequent trials. We then computed how likely each model was to produce the simulated choice behaviour from all other models, including its own choice behaviour. We were able to recover each model's behaviour successfully. The entries in **d** show each model's posterior probability of generating all simulated choice data sets, assuming a uniform prior. All models were by far the most likely to produce their own choice data.

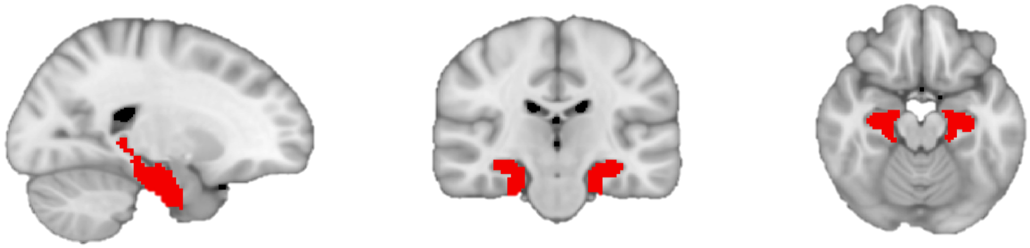

**Supplementary Figure S5. Anatomically defined region of interest used for small-volume correction.** The mask comprises the bilateral hippocampus, entorhinal cortex and subiculum

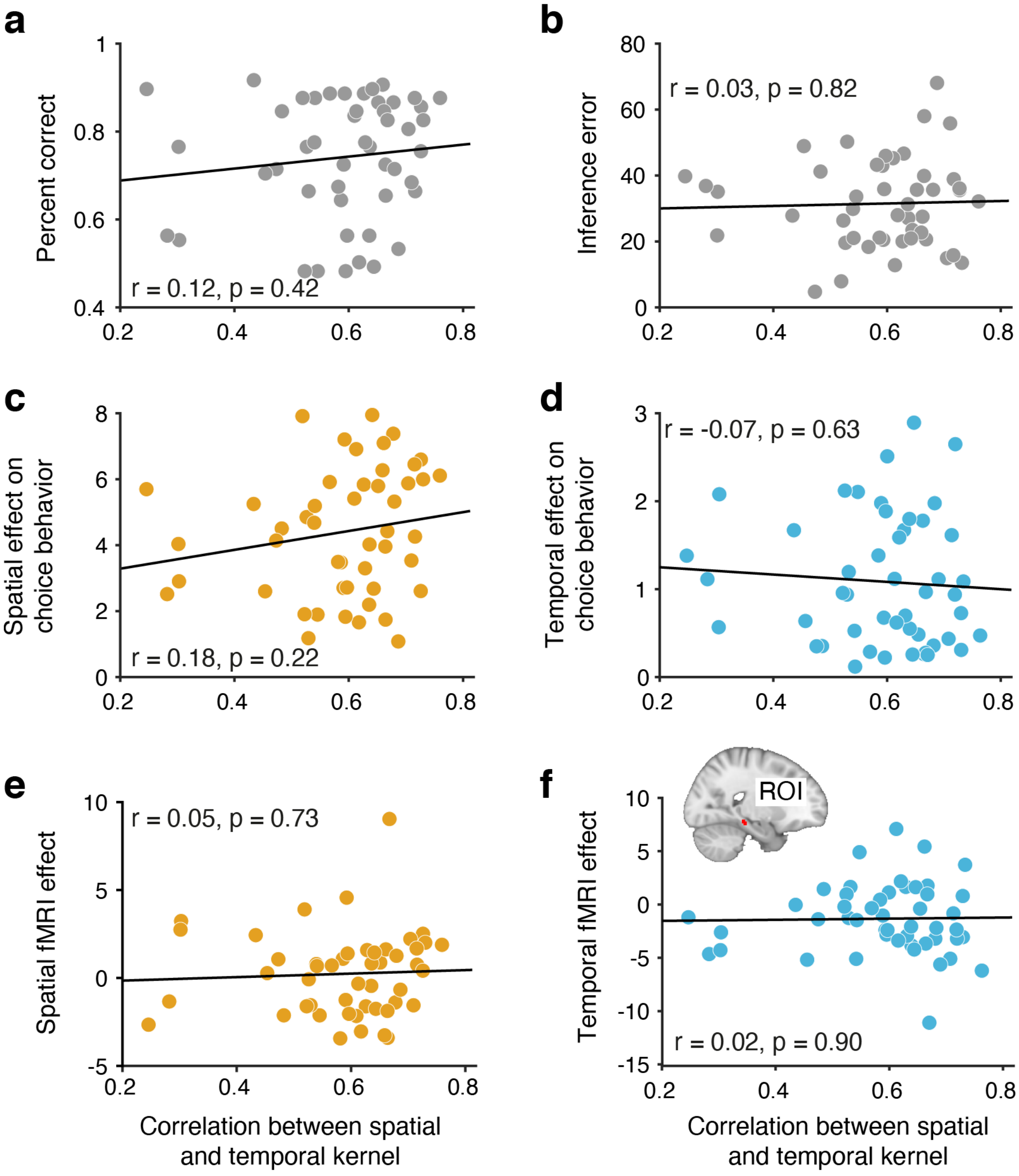


**Supplementary Figure S6. The correlation between spatial and the temporal kernels is not related to behavioral performance measures or hippocampal map representations**. The correlation between the spatial and the temporal kernel is plotted against and percent correct in the choice task (**a**), inference error(**b**), spatial effect on choice behavior (**c**), temporal effect on choice behavior (**d**) and fMRI cross-stimulus enhancement effect in the hippocampus for spatial (**e**) and temporal distances (**f**). Parameter estimates in **e** and **f** are extracted from the region of interest depicted in Figure 4a. None of the correlations reach significance (all $p > p0.2$).

### Supplementary Methods

#### Deriving the temporal kernel

Given a participant's exploration run from day 1, we want a method for obtaining a transition matrix $\mathbf{T}(s, s')$, whose entries reflect the participant's propensity for venturing directly from stimuli $s$ to stimuli $s'$. The successor representation (SR) is captured in the matrix $\mathbf{M}$, where entries $\mathbf{M}(s, s')$ equal the expected discounted number of future visits to stimulus $s'$, starting from $s$. If we know the transition matrix $\mathbf{T}$ governing the one-step transition probabilities between every pair of stimuli, we can define the SR matrix $\mathbf{M}$ as the following infinite sum of $\mathbf{T}$ raised to the power of $t$

$$\mathbf{M} = \sum_{t=0}^{\infty} \gamma^t \mathbf{T}^t \tag{7}$$

where $\gamma$ is the discount factor. This infinite sum can be computed analytically with matrix inversion

$$\mathbf{M} = (\mathbf{I} - \gamma \mathbf{T})^{-1} \tag{8}$$

where $\mathbf{I}$ is the identity matrix. Since we can compute the SR matrix $\mathbf{M}$ analytically from the transition matrix $\mathbf{T}$, we can attempt to recover the transition matrix from the SR matrix. This is fairly simple using matrix algebra. Since taking the matrix inverse of an inverted matrix gives us the uninverted matrix, $\mathbf{A}^{-1^{-1}} = \mathbf{A}$, we obtain

$$\mathbf{M}^{-1} = (\mathbf{I} - \gamma \mathbf{T}) \tag{9}$$

From Equation 9 we obtain $\mathbf{T}$ by subtracting the identity matrix $\mathbf{I}$, and dividing by $-\gamma$.

$$\mathbf{M}^{-1} - \mathbf{I} = -\gamma \mathbf{T} \tag{10}$$

$$\frac{\mathbf{M}^{-1} - \mathbf{I}}{-\gamma} = \mathbf{T} \tag{11}$$

leaving us with the transition matrix $\mathbf{T}$, which is such that performing an infinite random walk on it produces the SR matrix asymptotically.

#### Temporal relations explain reward generalization in the choice task

We sought to verify that the particular exploration trajectory a participant took on day 1 actually influenced how that participant generalized about value, and that the predictive performance of the temporal and the spatio-temporal model could not be attributed to other, more general properties of the temporal kernels, for instance, that they are generally similar to the spatial kernel. To test this, we shuffled the assignments of the temporal kernels, so that each participant would have their choices predicted based on a kernel computed from an exploration trajectory they *themselves* had not taken. If participant choices and generalization were really driven by their specific temporal interaction with the stimuli, then the predictive performance of a model based on the shuffled kernels should be substantially worse than the performance of a model using the correct exploration trajectories. We made the assignments symmetric (for a select pair of participant, their temporal kernels were swapped), and unique (no two participants could be assigned the same temporal kernel). As can be observed in Figure S4a, there were several temporal kernels that were substantially correlated with each other. We reasoned that swapping correlated kernels would yield smaller differences in predictive performance. We therefore sought to generate our shuffled assignments so that the overall correlation would be as small as possible. To do this, we sampled new kernels for each participant based on their inverse correlation $r^{-1}$ to the participant's true kernel. We sampled from a distribution obtained through a softmax transform

$$p_i(K_j) = \frac{exp(r_{ij}^{-1}/\tau)}{\sum_j^M exp(r_{ij}^{-1}/\tau)} \tag{12}$$

where $K_j$ is the kernel of participant $j$, $M$ is the number of participant minus participant $i$ and those already assigned, and $\tau$ is the temperature parameter. $\tau$ plays a key role here, as it allows us to control the degree to which we sample exclusively from the least correlated kernels, as opposed to more uniformly from all other kernels. As $\tau$ increases, the distribution gets more uniform. We collected negative log-likelihoods from the temporal model predicting participant choices (Figure S4b).

We sampled 10 kernel assignments for 10 evenly spaced values for $\tau$ between 0.01 and 3, leaving us with 100 samples of shuffled assignments, where the assignments were d with various degrees of uniformity. Substantiating the hypothesis that participant-specific temporal relations guide generalization, we observe that for all values of $\tau$, the shuffled assignments (the red line) produce substantially worse fits (negative log-likelihood) to the choice data on average than the model using each participant's true temporal kernel (dashed blue line). Moreover, we observe that this loss is at its highest on average when we sample kernels more concentrated based on inverse correlations (lower $\tau$), as opposed to more uniformly (higher $\tau$) from the set of all kernels.

#### Hyper-parameters

The successor representation was learnt with temporal-difference learning, using a discount rate $\gamma$ of 0.9. The signal variance parameter $\sigma_f^2$ of the Gaussian kernel (Equation 2) was set to 1, and the observation noise parameter $\sigma^2$ (Equation 1) was set to 0.01. The lengthscale parameter $\lambda$ of the diffusion kernel (Equation 6) was set to 1. For the spatial model, the best-fitting lengthscale $\lambda$ was 1.242. For the temporal model, the best-fitting learning rate $\eta$ was 0.4125. For the spatio-temporal model, the best-fitting lenghtscale $\lambda$ was 2.05, and the best-fitting learning rate $\eta$ was 0.01. To create the kernel matrices used as predictors in the fMRI analyses, we used the spatial kernel with a lengthscale of 2.05 and the temporal kernel with a learning rate of 0.01, which gave the best fit for the spatio-temporal model. These best-fitting hyper-parameter configurations were used in modelling value ratings, and for the model recovery.
